# Self-stabilization mechanism encoded by a bacterial toxin facilitates reproductive parasitism

**DOI:** 10.1101/2023.02.08.527603

**Authors:** Toshiyuki Harumoto

## Abstract

A wide variety of bacterial endosymbionts in insects are associated with reproductive parasitism, whereby they interfere with host reproductive systems to spread within populations. Recent successes in identifying bacterial factors responsible for reproductive parasitism have highlighted the common appearance of deubiquitinase domains, although their functional roles remain unknown. For example, *Spiroplasma* symbionts in *Drosophila* selectively kill male progeny with a male-killing toxin Spaid that encodes an OTU deubiquitinase domain. Here I show that without the function of OTU, the male-killing activity of Spaid is attenuated, though not eliminated, since it is polyubiquitinated and degraded through the host ubiquitin-proteasome pathway. Furthermore, I find that Spaid utilizes its OTU domain to deubiquitinate itself in an intermolecular manner. Collectively, the deubiquitinase domain of Spaid serves as a self-stabilization mechanism to facilitate male killing in flies, optimizing a molecular strategy of endosymbionts that enables the efficient manipulation of the host at low-cost.

## INTRODUCTION

Bacterial endosymbionts in insects and many other arthropods are exclusively transmitted to the next generation through the female germline (maternal transmission). Thus, endosymbionts face dead ends in males, no longer committed to expand their endosymbiont infection within host populations. To counter such a disadvantage, some endosymbionts manipulate host reproduction to increase the ratio of infected females by various means. These extraordinary phenomena, known collectively as reproductive parasitism, include feminization (conversion of genetic males into fertile females), parthenogenesis (enabling reproduction without mating with males), cytoplasmic incompatibility (CI) (killing of all progeny from matings between uninfected females and infected males), and male killing (selective killing of male progeny) (*1, 2*). Among them, male killing has evolved independently in distinct taxa of bacteria, such as *Wolbachia, Rickettsia*, and *Spiroplasma*, which are hosted by a variety of insect species having diverse sex-determination/differentiation systems (*3*).

To date, the underlying molecular mechanism of male killing is well studied in *Spiroplasma*-*Drosophila* symbiosis (*4, 5*). *Spiroplasma poulsonii* is a helical and motile bacterial endosymbiont that naturally inhabits *Drosophila* species such as *D. melanogaster* (*6*–*8*). Recently, a long-sought causative agent of male killing (i.e. male-killing toxin) in *D. melanogaster* was identified in the *S. poulsonii* genome and designated Spaid (*S. poulsonii* androcidin) (*9*). Its artificial expression in flies reproduces male-killing-associated pathologies that include abnormal apoptosis and neural defects during embryogenesis (*9*–*14*); moreover, green fluorescent protein (GFP)-tagged Spaid was observed to highly accumulate on the male X chromosome labeled with the dosage compensation machinery (*15*), congruent with the DNA damage/chromatin bridge-breakage specifically induced upon that chromosome (*9, 16*–*18*) (see also Fig. 3 and S3).

The amino acid sequence of Spaid encodes two eukaryotic-like domains (*9*) (Fig. 1). One is the ankyrin repeat, a common motif for protein-protein interactions, found not only in eukaryotic proteins but in proteins from pathogenic or symbiotic bacteria, archaea, and viruses (*19, 20*). Spaid without ankyrin repeats fails to accumulate on the male X chromosome and loses its male-killing activity, suggesting the ankyrin repeats play a pivotal role in targeting Spaid to the male X chromosome (*9*). The other domain is an ovarian tumor (OTU) domain, which encodes a deubiquitinase activity that reverses protein ubiquitination (*21*). Ubiquitination is an essential post-translational modification in eukaryotes, virtually affecting every aspect of proteins, such as stability, interactions, activity, and localization (*22*–*24*). In host-microorganism interactions, protein ubiquitination is crucial as a host defense against invading pathogens; conversely, a number of pathogenic bacteria and viruses employ deubiquitinase domain-containing proteins to counteract host ubiquitination machinery, and further exploit it for their own sake (*25*–*28*). OTU is one of seven deubiquitinase family members in eukaryotes; meanwhile, a plethora of viruses as well as a few pathogenic bacteria, including *Chlamydia pneumoniae* and *Legionella pneumophila*, also encode OTU domains for immune evasion (*21, 27, 29*–*34*). Recent omics approaches have shed light on the possible common usage of the OTU deubiquitinase domain in reproductive parasites (*35*–*38*); however, their biological roles, in particular how the encoded deubiquitinase activity is associated with the manipulative phenotypes is poorly defined. Previously, I showed that the abundance of Spaid protein lacking the OTU domain is dramatically reduced within the nuclei of host cells, although it retains residual male-killing activity (*9*) (see also Fig. S2 and S3). Here, I perform *in vivo* and *in vitro* assays using a variety of Spaid derivatives and characterize the role of the OTU deubiquitinase as a self-stabilization mechanism that enables efficient reproductive parasitism in insect endosymbiosis.

**Fig. 1.**
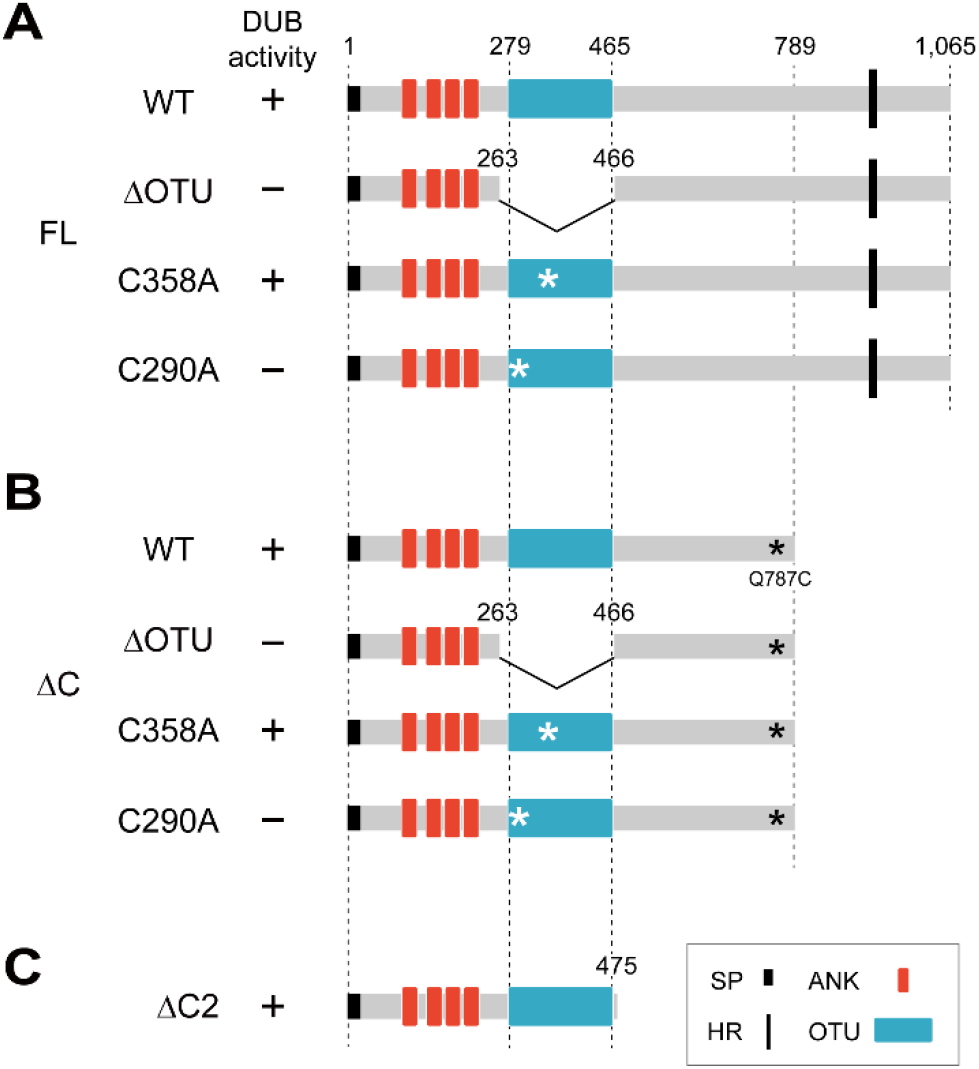
Structure of Spaid derivatives. Schematic representation of the protein structure of FL (**A**), ΔC (**B**), and ΔC2 Spaid (**C**). An N-terminal signal peptide (SP), ankyrin repeats (ANK, red), the OTU deubiquitinase domain (OTU, blue), and a C-terminal hydrophobic region (HR) are depicted as shown in the box. For the FL and ΔC forms, four derivatives including wild type (WT), lacking the OTU (ΔOTU), substituting presumed non-catalytic and catalytic cysteines into alanine (C358A and C290A, respectively) were generated. Asterisks represent the positions of amino acid substitutions. Presumptive deubiquitinase (DUB) activity of each derivative is indicated by “+” (active) and “–” (inactive). See text for other explanations.

## RESULTS

### The OTU deubiquitinase domain predicted in Spaid

Full-length Spaid (FL Spaid) is a 1,065 amino acid (aa) protein (Fig. 1A). Conserved domain searches by InterPro (*39*) predicted the OTU domain, spanning residues 279-465, in the Spaid aa sequence (Fig. 1A). The OTU family proteins are cysteine proteases, whose catalytic Cys-His dyad constitutes the active site of the deubiquitinating enzyme (*21, 27*). The multiple alignment of the predicted OTU in Spaid with representative family members from both eukaryotes and bacteria revealed putative catalytic residues Cys-290 and His-458 in the OTU sequence of Spaid (Fig. S1A). I further scrutinized the above domain annotation by applying two distinct methods of protein structure modeling to the presumptive Spaid OTU sequence. First, I utilized the Phyre2 homology modeling server (*40*), and found that the high-ranked 3D fold models (with confidence values of > 90%) are all generated using the OTU family proteins as templates (Table S1, sheet1). Next, I employed ColabFold software (*41*) whose structural prediction is powered by the AlphaFold2 program, a neural network model integrating physical and biological features of protein structure (*42*). Structural comparison of the ColabFold model against the Protein Data Bank (PDB) by a DALI server search (*43*) successfully retrieved OTU family proteins as top hits (with Z scores of 6.2-14.1) (Table S1, sheet2). In the above two 3D models, Cys-290 and His-458 near the N- and C-terminal domain boundaries were in close vicinity to each other, which was very likely to constitute the catalytic Cys-His dyad for the Spaid OTU (Fig. S1, B and C). These data, together with the following genetic/biochemical analyses of a catalytic Cys-290 mutant (see below), reinforce the idea that Spaid possesses deubiquitinase activity through its OTU domain.

To address the function of the Spaid OTU in male killing, I previously generated a deletion construct lacking residues 264-465, encompassing the OTU domain (ΔOTU, Fig. 1A). In this study,, I produced an amino acid substitution construct by replacing Cys-290 with alanine, expecting to abolish deubiquitinase activity (C290A, Fig. 1A). I also substituted another presumed non-catalytic cysteine (Cys-358) to serve as a negative control (C358A, Fig. 1A). I utilized these four Spaid derivative constructs (WT and C358A, deubiquitinase active controls; ΔOTU and C290A, presumptive deubiquitinase inactive derivatives; Fig. 1A) to confirm the deubiquitinase activity of Spaid and explore its exact role in male killing.

### C290A mimics the phenotypes of the OTU deletion

In my previous paper, I employed the GAL4/UAS system to express WT and ΔOTU in *D. melanogaster* as GFP-tagged proteins and compared their male-killing activity and subcellular localization patterns (*9*). I repeated the experiments and reproduced the results as follows: i) strong expression of WT and ΔOTU by the *actin-GAL4* ubiquitous driver eliminated male progeny regardless of the presence or absence of the OTU domain (Fig. S2A); ii) alternatively, when weakly expressed by the *armadillo-GAL4* driver, WT Spaid still killed all males, but the ΔOTU construct could no longer kill them (Fig. S2B). A re-examination of the subcellular distribution of GFP-tagged WT and ΔOTU Spaid in larval salivary gland cells confirmed their distribution in the cytoplasm and/or more likely in the intracellular membrane system including the endoplasmic reticulum (ER), Golgi apparatus, and plasma membrane irrespective of sexes (Fig. S3, A and B; see also the illustration in Fig. S4A). A striking difference was evident in the nuclear localization of Spaid-GFP in cells, as shown previously, where the ΔOTU Spaid-GFP signals were reduced (Fig. S3, A and B); additionally, the strong accumulation on the male X chromosome and resultant DNA damage were greatly attenuated by the deletion of the OTU domain (Fig. S3, B and C).

I then repeated the above experiments with the newly designed C358A and C290A constructs and obtained comparable results: i) strong expression of both constructs eliminated male offspring completely (Fig. S2A), while weak expression of C358A but not C290A showed a substantial male killing phenotype (Fig. S2B), ii) the distribution patterns of C358A and C290A were almost the same as those of WT and ΔOTU, respectively (Fig. S3). Thus, a single amino acid mutation at C290 closely mimics the phenotypes observed with the entire deletion of the OTU domain, confirming C290 as a key residue in the active site of the OTU domain, which works as a cysteine-type deubiquitinase. The data also suggest that the OTU deubiquitinase activity is not essential for the male-killing activity itself. Rather, it is assumed to be involved in the nuclear translocation of the protein or the enhancement of protein stability to facilitate male killing activity.

### Loss of OTU function reduces Spaid protein levels

During my initial characterization of the *spaid* locus, I obtained a spontaneously mutated *Spiroplasma* strain whose male-killing ability is partially lost (*9*). Comparison of the genomic sequence with that of the original strain revealed an 828-bp deletion, leading to a C-terminally truncated form of Spaid (ΔC Spaid) lacking the hydrophobic region (HR) (Fig. 1B) (*9*). In the same manner as for FL Spaid, I constructed and expressed WT, ΔOTU, C358A, and C290A derivatives of ΔC Spaid (Fig. 1B) with strong/weak GAL4 drivers and obtained results similar to those with FL Spaid (Fig. 2), indicating that the HR of FL Spaid is dispensable for the male killing activity under the artificial expression condition, although its loss in *Spiroplasma* reduces male-killing ability (*9*). The exact function of the HR remains unknown, but it may affect the behavior of Spaid within bacterial cells, potentially in membrane targeting and/or extracellular secretion.

**Fig. 2.**
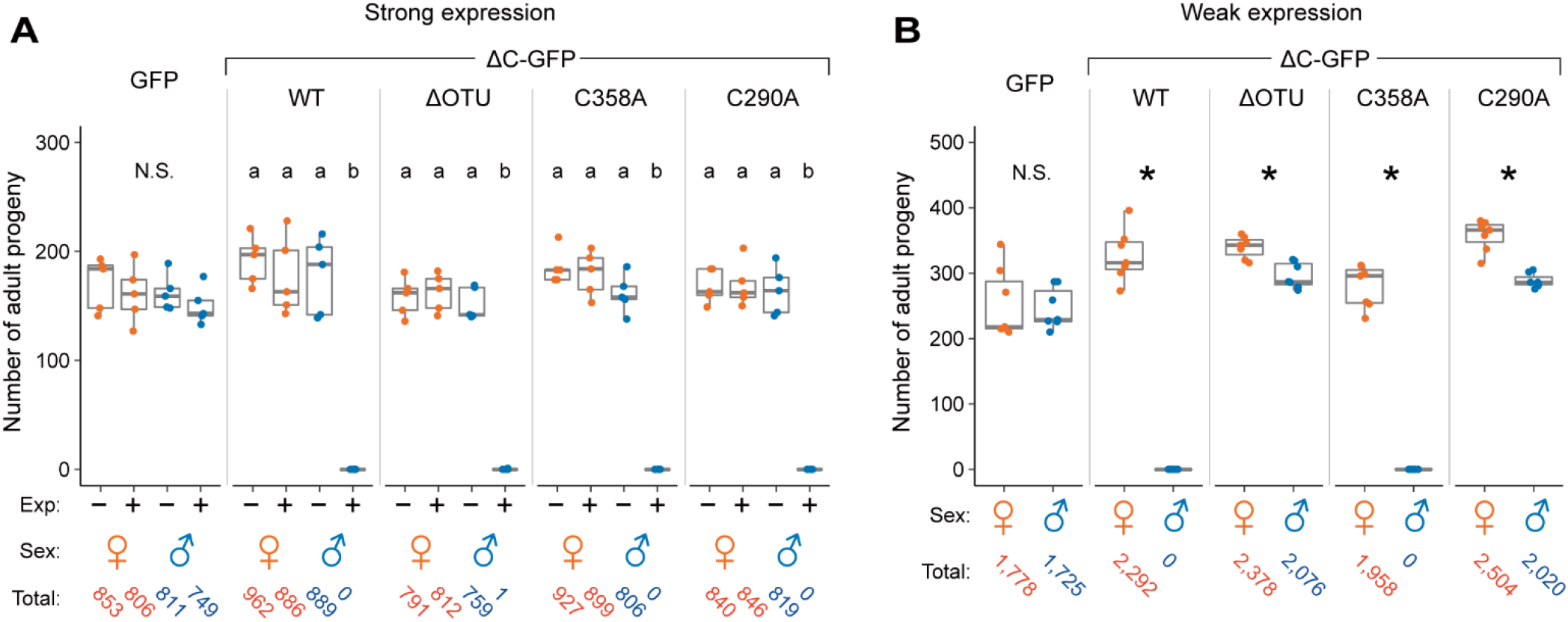
The OTU-deficiency attenuates the male-killing activity of ΔC Spaid. *UAS-GFP* (negative control), *UAS-ΔC Spaid* (WT) and its derivative lines (ΔOTU, C358A and C290A) were crossed with the *actin-GAL4* (**A**, strong expression; n = 5 independent crosses) and *armadillo-GAL4* (**B**, weak expression; n = 7 independent crosses) driver lines, respectively. The numbers of female (red) and male (blue) progeny obtained from the crosses are shown. In **A**, offspring with both *GAL4* and *UAS* (+) or with only *UAS* (–) were counted separately. Different letters (Steel–Dwass test, **A**) and asterisks (two-tailed Mann–Whitney U test, **B**) represent statistically significant differences (P < 0.05). NS, not significant (P > 0.05) (See also Table S2). Box plots indicate the median (bold line), upper and lower quartiles (box edges), and the maximum/minimum values (whiskers). Dot plots show individual data points. The total counts for each genotype and sex are shown at the bottom.

Next, I examined the subcellular localization pattern of ΔC Spaid in larval salivary gland cells by monitoring GFP-tagged constructs. There were marked differences. Unlike FL Spaid, the GFP signals of WT and C358A ΔC Spaid were predominantly detected within nuclei but were negligible in the intracellular membrane system (Fig. 3, A and B; see also the illustration in Fig. S4B). Most strikingly, the GFP signals of ΔOTU and C290A ΔC Spaid were extremely diminished within the cells, and the intranuclear signals were far more affected in both sexes (Fig. 3). The distinct distribution patterns of the FL and ΔC forms indicated that the HR likely dispatches Spaid to the host intracellular membrane system when artificially expressed by the GAL4/UAS system (Fig. S4).

**Fig. 3.**
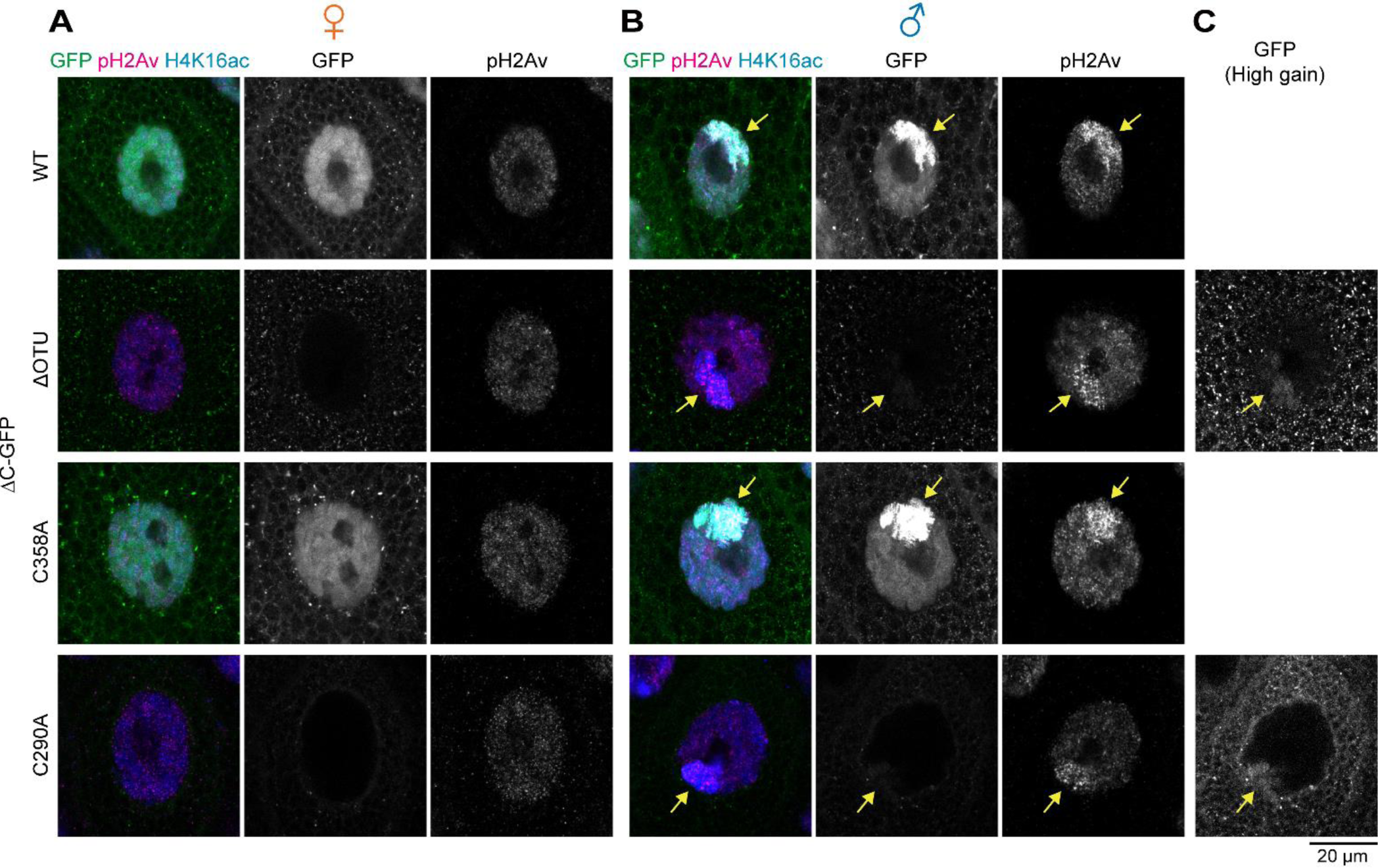
OTU-deficiency reduces the expression level of ΔC Spaid. Female (**A**) and male (**B**) larval salivary gland cells expressing ΔC Spaid-GFP (green). WT (female, n = 11; male, n = 8), ΔOTU (female, n = 13; male, n = 8), C358A (female, n = 14; male, n = 9), and C290A (female, n = 12; male, n = 12) derivatives were immunostained for DNA damage (pH2Av, magenta) and histone H4 lysine 16 acetylation (H4K16ac, blue). Yellow arrows represent the accumulation of ΔC Spaid-GFP and pH2Av on the male X chromosome labeled with the dosage compensation machinery (marked with strong H4K16ac signals). (**C**) High-gain images of ΔOTU and C290A reveal faint GFP signals on the male X chromosome.

Further, these results strongly suggested that the OTU domain is not involved in the nuclear translocation of Spaid, but rather is required for its stable expression. In the following experiments, I tested this possibility by using the ΔC forms of Spaid, whose distribution patterns seemed to be more amenable to the analysis of protein stability.

As expected, WT/C358A forms of ΔC Spaid strongly accumulated on the male X chromosome and induced DNA damage, similar to FL Spaid (Fig. 3B). In contrast, the ΔOTU and C290A forms of ΔC, but not FL, Spaid induced DNA damage on the male X chromosome to some extent (compare Fig. 3, B and C with S3, B and C), probably due to the higher accumulation of ΔC Spaid inside the nucleus, hence on the male X chromosome. These results also provided molecular evidence for the dispensability of the deubiquitinase activity of Spaid for the DNA damaging activity.

### The deubiquitinase activity of Spaid counteracts host proteasome-dependent degradation

To assess whether the low abundance of OTU-deficient Spaid (ΔOTU/C290A) is related to the ubiquitin-proteasome degradation pathway, I transiently expressed ΔC Spaid-GFP in *Drosophila* S2 cells and applied a proteasome inhibitor, MG-132, to block the host proteasomal activity. In the absence of MG-132, the levels of GFP fluorescence of OTU-deficient constructs were significantly lower than those of control constructs, reminiscent of the results in the larval salivary gland cells, whereas they were substantially recovered after the treatment with MG-132 (Fig. 4A and S5A).

**Fig. 4.**
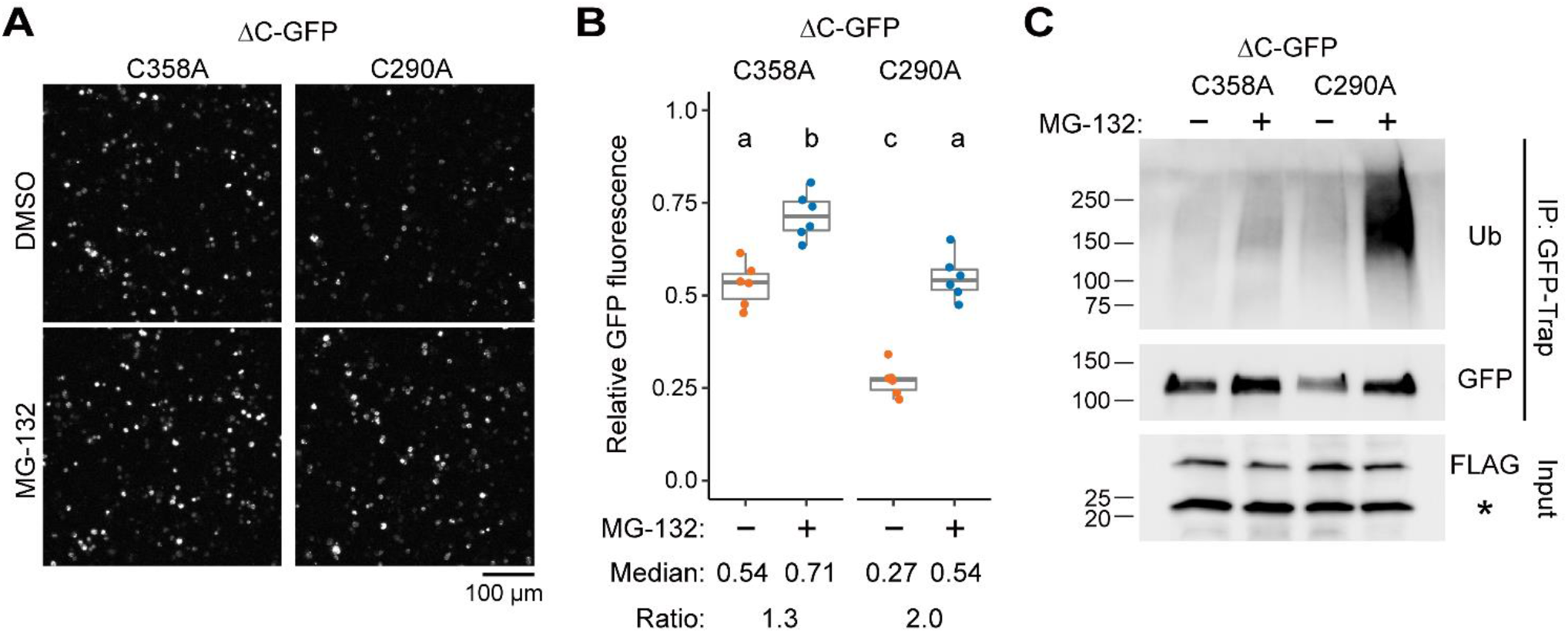
The OTU deubiquitinase of Spaid stabilizes itself by counteracting the host ubiquitin-proteasome pathway. (**A**) S2 cells expressing ΔC Spaid-GFP. The raw GFP signals of C358A and C290A derivatives are indicated. Cells were treated with DMSO (top) or proteasome inhibitor MG-132 (bottom) for 9 h. (**B**) The relative GFP fluorescence of C358A and C290A derivatives of ΔC Spaid-GFP expressed in S2 cells cultured without/with MG-132, respectively. The same data sets used in (**A**) were analyzed (n = 6 areas per well in each condition). As a transfection marker, FLAG-tagged mCherry (mCherry-FLAG) was co-transfected and its fluorescence was used to normalize the GFP fluorescence (see Materials and Methods). The median values and the ratios derived from GFP fluorescence in the absence and presence of MG-132 are shown at the bottom. Different letters (Steel–Dwass test) represent statistically significant differences (P < 0.05) (See also Table S2). (**C**) Ubiquitination assay using GFP-Trap. C358A and C290A derivatives of ΔC Spaid-GFP were immunoprecipitated (IP) from S2 cells cultured without/with MG-132. Immunoprecipitates were washed with stringent buffer and analyzed by Western blotting to detect GFP (represents native ΔC Spaid) and ubiquitin (Ub; a high-molecular-weight smear represents the polyubiquitinated fraction). Input samples were probed with anti-FLAG antibody. mCherry-FLAG and non-specific bands (“FLAG” and “*”) are shown as transfection and loading controls, respectively.

Quantification of the relative GFP fluorescence revealed that it is elevated with MG-132 treatment in general; however, the GFP fluorescence ratio between with/without MG-132 treatment was higher for OTU-deficient constructs than for control constructs (Fig. 4B and S5B), indicating that the stability of OTU-deficient Spaid is much more sensitive to the host proteasomal activity.

Then how does Spaid counteract host proteasomal degradation with the help of its OTU deubiquitinase activity? The simplest scenario is that Spaid can remove the attached ubiquitin conjugation by itself (auto-deubiquitination). In that case, OTU-deficient Spaid, which is polyubiquitinated and degraded in the absence of MG-132, would accumulate and be detected in the presence of MG-132. To test this directly, ΔC Spaid-GFP was expressed and precipitated from whole cell extracts with GFP-Trap magnetic beads, and bound proteins were isolated by highly stringent washes (8M Urea, 1% SDS in PBS; see Materials and methods) (*44, 45*). The precipitated ΔC Spaid-GFP was analyzed for the ubiquitination state using a ubiquitin-specific antibody for Western blot analysis. As expected, a high molecular weight smear was detected only in the samples expressing OTU-deficient constructs treated with MG-132, corresponding to the polyubiquitinated fraction of ΔC Spaid (Fig. 4C and S5C). No obvious polyubiquitinated signals were observed in OTU active control samples, biochemically confirming the deubiquitinase activity of Spaid OTU. These results together with the harsh conditions used in the immunoprecipitation experiments supported the idea that Spaid is auto-deubiquitinated to protect itself from proteolysis by the host proteosome.

### Self-stabilization of Spaid through intermolecular deubiquitination

Several mammalian deubiquitinases have been shown to deubiquitinate themselves to change their own ubiquitination states and resultant fates (*46*–*51*). More recently, a comprehensive library screening identified a set of mammalian deubiquitinases that auto-deubiquitinate themselves either in an intramolecular (within the protein) or intermolecular (between the proteins) manner to increase their own stability (*52*). To better understand the behavior of Spaid OTU deubiquitination, I sought to determine which strategy is applicable to Spaid. To this end, I designed co-transfection experiments with two differently tagged ΔC Spaid constructs with/without deubiquitinase activity.

C290A-GFP (deubiquitinase-defective) expressing cells were co-transfected with either a C358A-HA (deubiquitinase-proficient) or a C290A-HA construct. I first analyzed the expression level of C290A-GFP and found that the GFP fluorescence was relatively higher in the co-transfection with C385A-HA than with C290A-HA, even without the addition of MG-132 (Fig. 5, A and B), implying that the proteolysis of C290A is blocked in the presence of C358A. Next, C290A-GFP was precipitated from the respective whole cell extracts in denaturing conditions, and its polyubiquitination state was analyzed with the same technique as for the ubiquitination assay.

**Fig. 5.**
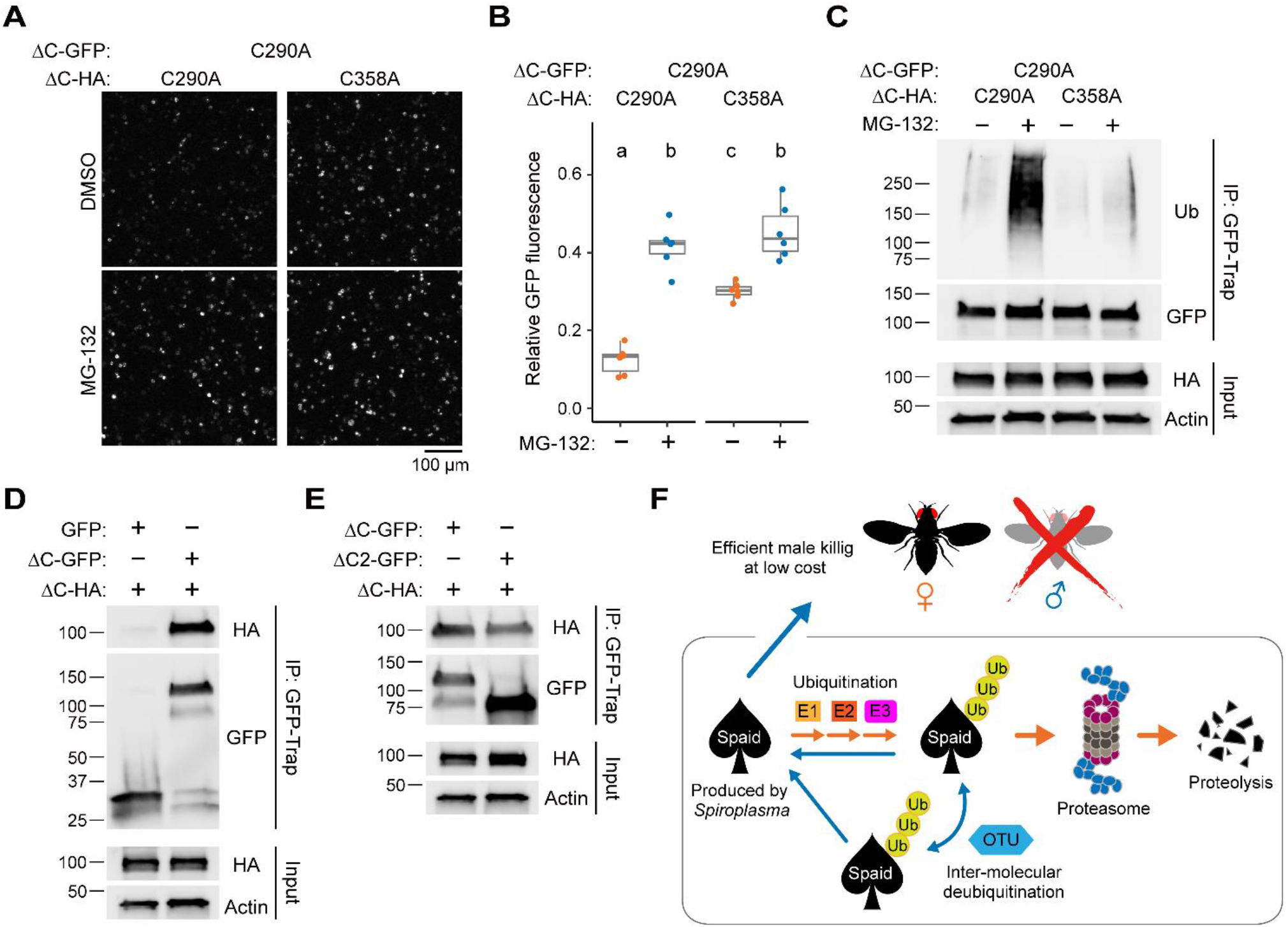
Self-stabilization of Spaid by intermolecular deubiquitination. (**A**) S2 cells expressing the C290A derivative of ΔC Spaid-GFP, co-transfected with a C290A or C358A derivative tagged with HA, respectively. The raw GFP signals of C290A are indicated. Cells were treated with DMSO (top) or MG-132 (bottom) for 9 h. (**B**) The relative GFP fluorescence of the C290A derivative of ΔC Spaid-GFP, co-transfected with a C290A or C358A derivative tagged with HA in S2 cells cultured without or with MG-132. The same data sets used in (**A**) were analyzed (n = 6 areas per well in each condition). Other explanations are the same as in Fig. 4B. Different letters (Steel–Dwass test) represent statistically significant differences (P < 0.05) (See also Table S2). (**C**) Ubiquitination assay using GFP-Trap. The same technique used in Fig. 4C was applied. Input samples were immunoblotted and probed with anti-HA and anti-β-Actin antibodies to show the transfection efficiency and the loading control. (**D**) Homomeric interaction of ΔC Spaid in S2 cells. ΔC Spaid-GFP was co-immunoprecipitated with ΔC Spaid-HA. Free GFP was used as a negative control. Input samples were immunoblotted with anti-HA and anti-β-Actin antibodies to show the transfection efficiency and the loading control. (**E**) Direct interaction of ΔC and ΔC2 Spaid in S2 cells. ΔC2 Spaid-GFP was co-immunoprecipitated with ΔC Spaid-HA. ΔC Spaid-GFP was used as a positive control. Input samples were immunoblotted with anti-HA and anti-β-Actin antibodies to show the transfection efficiency and the loading control. (**F**) A proposed model for the self-stabilization mechanism of Spaid to facilitate male killing in flies. See text for details.

Remarkably, polyubiquitination of C290A-GFP disappeared when the cells were co-transfected with C358A-HA, but was not affected by co-transfection with C290A-HA (Fig. 5C). These data support the notion that the auto-deubiquitination of Spaid occurs in an intermolecular manner, although they did not necessarily exclude the possibility that some intramolecular deubiquitination can occur.

If intermolecular auto-deubiquitination of Spaid occurs, a direct homomeric interaction of Spaid would be expected. To test this, cells were co-transfected with ΔC Spaid-GFP and ΔC Spaid-HA, and their direct interaction was analyzed by a co-immunoprecipitation assay. Cells co-transfected with GFP and ΔC Spaid-HA served as a negative control. Western blot analyses of GFP-Trap precipitates with an anti-HA antibody showed that the HA signal was detected from the ΔC Spaid-GFP and ΔC Spaid-HA co-transfected co-immunoprecipitation, but not from the negative control, demonstrating the homomeric interaction of Spaid *in vivo* (Fig. 5D).

Finally, I tried to narrow down the responsible amino acid region for the homomeric interaction of Spaid. The two eukaryotic-like domains encoded in the N-terminal half of Spaid, ankyrin repeats and the OTU domain, are involved in targeting Spaid to the male X chromosome (*9*) and protein self-stabilization (this study), respectively. Therefore, I suspected that the remaining C-terminal half without any functional prediction could be involved (Fig. 1). I generated a construct lacking almost the entire sequence following the OTU domain (ΔC2 Spaid; Fig. 1C) and performed a co-immunoprecipitation assay. Similar to ΔC Spaid-GFP, ΔC2 Spaid-GFP was also precipitated together with ΔC Spaid-HA (Fig. 5E), thus excluding the simple possibility that the C-terminal half of the protein is solely required for the homomeric interaction.

## DISCUSSION

In this study, I characterized the role of the OTU deubiquitinase domain of Spaid in male killing. I have shown that: i) the male-killing toxicity of Spaid is attenuated but not eliminated without a functional OTU; ii) the OTU-deficient Spaid is unstable within host cells, because it is polyubiquitinated and degraded through the host ubiquitin-proteasome system; iii) the OTU domain works intermolecularly to deubiquitinate Spaid; and iv) Spaid monomers directly interact to form homomers, though its C-terminal region is dispensable for the interaction. Taken together, these data lead to a model in which the OTU deubiquitinase of Spaid serves as a self-stabilization mechanism to facilitate male killing in flies (Fig. 5F). In my artificial expression experiments using the weak GAL4 driver (*armadillo-GAL4*), the OTU-deficient Spaid failed to kill males (Fig. 2B and S2B). Assuming that even the weak GAL4 driver would produce much more Spaid than the endogenous bacteria within host cells, increasing protein stability through the OTU deubiquitinase domain is indispensable for the efficient killing of male flies in the natural symbiotic relationship (Fig. 5F). Given that the available resources are limited in endosymbiosis, the presence of a self-stabilization mechanism, at zero energy cost to the bacteria, would be fundamental for reproductive parasitism.

In my previous study (*9*), I hypothesized that the OTU domain of Spaid is involved in nuclear translocation, based on the localization analysis of FL Spaid and its deletion constructs (see also Fig. S3). The present study using ΔC Spaid and derivatives describes the deubiquitinase activity of OTU and its biological role in evading the host ubiquitin-proteasome degradation. In general, active proteasomes are ubiquitously present in the nucleus, cytoplasm, the cytoplasmic side of membranous organelles including the ER, but not inside the lumen (*53*). This may account for the disappearance of OTU deficient FL Spaid only from the nucleus as follows: OTU-deficient Spaid is degraded by proteasomes within the nucleus and cytoplasm; however, the fraction targeted to the intracellular membrane system with the help of HR (see also Fig. S4) is not exposed to active proteasomes for degradation.

A number of bacterial deubiquitinases, like OTU, are distantly related members of eukaryotic deubiquitinase families, although most of them belong to a distinct cysteine peptidase family of a different origin, categorized as the clan CE (*27, 54, 55*). One of the best characterized proteins containing a CE peptidase domain in reproductive parasites is a *Wolbachia* factor responsible for CI (*1*). CI factors (Cifs) consist of a binary gene product of CifA and CifB, and the latter contains the Ulp1 (ubiquitin-like-specific protease 1) domain whose deubiquitinase activity is biochemically validated (*56*–*58*). A recent *in vivo* expression study of CifB in *D. melanogaster* suggested that its deubiquitinase activity could contribute to self-stabilization within host cells (more precisely, spermatid nuclei). Without the intrinsic deubiquitinase activity, its protein level in spermatids declined, whereas high-level expression is normally required for the induction of male sterility (*59*). Very recently, a novel *Wolbachia* protein designated Oscar was identified as a causative factor of male killing in Lepidoptera (*60*). The authors showed that the expression of Oscar kills males by directly targeting and degrading host Masculinizer protein required for masculinization and dosage compensation (*60*–*62*). Interestingly, Oscar contains a CifB-like domain, possibly possessing Ulp1 enzymatic activity, yet it does not seem to be essential for the male-killing function of Oscar (*60*).

Although detailed molecular validation is still awaited, these two *Wolbachia* proteins dealing with distinct reproductive manipulations, CI and male killing, might commonly employ their deubiquitinase domains for the stable expression within host cells/tissues, analogous to the Spaid OTU.

The present study raises further questions. First, the identity of the host molecules and pathways involved in the ubiquitination/degradation of Spaid are unknown. Protein ubiquitination is catalyzed by a cascade of E1, E2, and E3 enzymes, and E3s are responsible for the substrate recognition and specificity (*22, 63*). It would be useful to further investigate whether a specific E3 ligase for Spaid is dispatched, or whether other conventional E3s that can recognize analogous host protein sequence motifs are applied for the process. In either case, the nuclear proteasome system could participate in the degradation, in agreement with the high sensitivity within nuclei without the function of OTU (Fig. 3 and S3). Second, the extent to which deubiquitinase-dependent stabilization mechanisms operate in other types of reproductive parasitism, including feminization and parthenogenesis, is unclear, because our knowledge about the molecular aspects of reproductive parasitism is still very limited: only factors responsible for male killing and CI in dipteran and lepidopteran host species have thus far been identified (*9, 56*–*58, 60, 64*). Third, the origin and evolution of these bacterial toxin/effector proteins is obscure. Multiple lines of evidence have suggested that the bacterial deubiquitinases were acquired from eukaryotic genomes via lateral gene transfer (*27, 36*). It is possible that the horizontally acquired deubiquitinase-containing locus was used as a scaffold for the assembly of toxin/effector proteins (*36*). Alternatively, the deubiquitinase domain could have been acquired afterwards as the result of an evolutionary arms race between hosts and bacterial endosymbionts. Thus, further studies are needed to provide an overall picture of reproductive parasitism in arthropods, which would be beneficial in developing innovative approaches to manipulate and control arthropods that could play both beneficial and harmful roles in natural and agricultural settings.

## MATERIALS AND METHODS

### Fly stocks and genetics

The stocks of *D. melanogaster* were reared at 25 °C with standard cornmeal food. The absence of *Spiroplasma* and *Wolbachia* were tested by a diagnostic PCR with specific primers (SpoulF 5’-GCT TAA CTC CAG TTC GCC-3’ and SpoulR 5’-CCT GTC TCA ATG TTA ACC TC-3’ for *Spiroplasma*; *wsp*_81F 5’-TGG TCC AAT AAG TGA TGA AGA AAC-3’and *wsp*_691R 5’-AAA AAT TAA ACG CTA CTC CA-3’ for *Wolbachia*) (*6, 65*). Some of them were treated with tetracycline before the test (*9*). The following lines were provided by the Bloomington *Drosophila* Stock Center at Indiana University (BDSC) and the Department of *Drosophila* Genomics and Genetic Resources at Kyoto Institute of Technology (DGGR): *actin-GAL4* (BDSC #4414), *tubulin-GAL80*^*ts*^ (BDSC #7108), *armadillo-GAL4* (BDSC #1560), *UASp-EGFP* (DGGR #116071), and *CyO, ActGFP* (DGGR #107783). *UASp-Spaid-EGFP* and *UASp-Spaid*.*ΔOTU-EGFP* transgenic lines were generated in the previous study (*9*). Other original *UAS* transgenic lines listed below were generated using germline transformation by P-element based plasmid vectors (BestGene Inc.): *UASp-Spaid*.*C290A-EGFP, UASp-Spaid*.*C358A-EGFP, UASp-ΔC Spaid-EGFP, UASp-ΔC Spaid*.*ΔOTU-EGFP, UASp-ΔC Spaid*.*C290A-EGFP*, and *UASp-ΔC Spaid*.*C358A-EGFP*. For the plasmid construction, please see below.

To artificially express *spaid* in flies by the GAL4/UAS system (*66*), homozygous *UAS-Spaid* transgenic lines listed above were crossed with the *actin-GAL4*/*CyO* (for the strong expression in Fig. 2A and S2A) or *armadillo-GAL4* (for the weak expression in Fig. 2B and S2B) driver lines. Emerging adult flies were segregated by sexes and genotypes (in the crosses with *actin-GAL4*/*CyO*) and counted separately until the 18-19th day in each vial. In the latter crosses with the *armadillo-GAL4* driver line, *UAS-Spaid* female flies were mated with GAL4 driver male flies to avoid the effect of maternally loaded GAL4 (*67*), which caused substantial male death during embryonic stages. In the former crosses with the *actin-GAL4* driver line, no obvious differences were observed in the reciprocal mating, suggesting no/negligible maternal contribution in the experiments. For the expression of *spaid* in larval salivary gland cells (Fig. 3 and S3), male lethality was circumvented by using a recombined *actin-GAL4, tubulin-GAL80*^*ts*^/*CyO* line. Crosses were maintained at 20 °C for 6-7 days to let the larvae grow until the early 3rd instar stage, then were shifted up to 29 °C and kept for 1 day before dissection at the wandering 3rd instar stage. Only GFP-positive larvae were dissected.

### Cell culture and transfection

*Drosophila* S2 cells (provided by Masayuki Miura) were maintained at 25 °C with Schneider’s *Drosophila* Medium (Gibco, 21720024) supplemented with Fetal Bovine Serum (Cytiva, SH30071.03) and penicillin-streptomycin (FUJIFILM Wako Pure Chemical, 168-23191). For plasmid DNA transfection, cells were seeded in multi-well plates (24-well plate with 500 µL medium for imaging; 6-well plate with 2 mL medium for biochemical assays) at 1.0 × 10^6^ cells/ml one day before transfection. The cells were transfected with pMT plasmid vectors (encoding ΔC Spaid derivatives tagged with GFP and mCherry as a transfection control; see below for details) using HilyMax transfection reagent (Dojindo, H357) following the manufacturer’s protocol. A few hours after transfection, the medium was changed to avoid toxicity. About 20 h after transfection, the metallothionein promoter was induced by adding 1mM CuSO4 to the medium. About 20 h after the induction, 20 µM MG-132 (Calbiochem, 474790) diluted in DMSO was added to inhibit the proteasomal activity. As a negative control, DMSO was added. 9 h after the treatment, the cells were observed or harvested for protein extraction.

### Plasmid construction

The codon-optimized *ΔC spaid* sequence (2,367 bp) was synthesized and cloned into the pDONR221 vector by GeneArt service (Thermo Fisher Scientific) in the previous study (*9*). The FL *spaid* sequence (3,195 bp) was generated and cloned into the pENTR vector as described previously (*9*).

To generate the ΔOTU deletion in ΔC Spaid, two PCR fragments (nucleotide positions 1-789 and 1,396-2,367)) were fused by PCR and cloned into the pENTR vector, using a pENTR/D-TOPO cloning kit (Thermo Fisher Scientific) as described previously (*9*). The C290A and C358A single amino acid substitution constructs were generated by the SPRINP (Single-Primer Reactions IN Parallel) method (*68*). Briefly, the template plasmid DNA (pENTR-FL Spaid and pDONR221-ΔC Spaid, respectively) was amplified with either the forward or reverse primer separately (C290A forward primer 5’-GGC TCC <u>GC</u>C CTG TTT TGG AGT GTG GCC-3’, C290A reverse primer 5’-AAA CAG G<u>GC</u> GGA GCC ATC CTC GAC CAC-3’; C358A forward primer 5’-GCC AAC <u>GC</u>C CTG ATC CGC GAT ATC TTC-3’, C358A reverse primer 5’-GAT CAG G<u>GC</u> GTT GGC GGT CTG ATC GCT-3; the underscores represent the nucleotide substitution introduced into the primers). The two single-primer PCR reactions were mixed and denatured, then cooled down gradually to reanneal the complementary strands. After purification with the Wizard SV Gel and PCR Clean-Up System (Promega), the template plasmid DNA was digested with DpnI and used in the transformation reaction.

UAS-Spaid plasmids generated for this study were as follows: pUASp-Spaid.C290A-EGFP; pUASp-Spaid.C358A-EGFP; pUASp-ΔC Spaid-EGFP; pUASp-ΔC Spaid.ΔOTU-EGFP; pUASp-ΔC Spaid.C290A-EGFP; pUASp-ΔC Spaid.C358A-EGFP; and pUASp-ΔC Spaid-3xHA. To construct these plasmids, the Gateway cassettes containing the *spaid* open reading frames (ORFs) in the pDONR221/pENTR vectors were transferred into the destination vectors pPWG (#1078) and pPWH (#1102), obtained from the *Drosophila* Genomics Resource Center (DGRC) by the LR clonase II enzyme mix kit (Thermo Fisher Scientific). *UAS-spaid* transgenic lines used in the fly genetics experiments (Fig. 2, 3, S2 and S3) were generated by microinjection of the above plasmids into *D. melanogaster* embryos (except for the pUASp-ΔC Spaid-3xHA plasmid used only for the vector construction below).

In the ubiquitination assay in S2 cells (Fig. 4, 5A to C, and S5), the following ΔC Spaid plasmids were used: pMT-ΔC Spaid-EGFP; pMT-ΔC Spaid.ΔOTU-EGFP; pMT-ΔC Spaid.C290A-EGFP; pMT-ΔC Spaid.358A-EGFP; pMT-ΔC Spaid.C290A-3xHA; and pMT-ΔC Spaid.358A-3xHA. To construct these plasmids, the Gateway cassettes containing *ΔC spaid* fragments in the pDONR221/pENTR vectors were transferred into the pMTWG destination vector containing the metallothionein promoter and a C-terminal GFP tag (a gift from Catherine Regnard and Peter Becker). To replace the EGFP tag with the 3xHA tag, a PCR fragment containing a 969-bp sequence of *ΔC spaid* (nucleotide position 1,399-2,367, containing an AgeI site) followed by the 3xHA sequence was PCR amplified from the pUASp-ΔC Spaid-3xHA plasmid (forward primer 5’-AAC ATC CGC ATG ATC AAC GAG-3’, reverse primer 5’-CTA <u>GCT AGC</u> TTA GTG TCC GCC ATG AGC AGC GTA ATC-3’; the underscore represents the NheI site added to the primer). Then, the AgeI-NheI fragment (1,079 bp) was inserted into the pMT-ΔC Spaid.C290A/C358A-EGFP plasmid, which had been digested with the corresponding restriction enzymes. To make a transfection control plasmid (pMT-mCherry-FLAG), the ORF of mCherry was amplified with primers containing restriction enzyme sites and the FLAG tag sequence (forward primer 5’-CCG <u>GAT ATC</u> CAA CAT GGT GAG CAA GGG CGA GGA G-3’, reverse primer 5’-CTA <u>GCT AGC</u> TTA CTT GTC ATC GTC GTC CTT GTA ATC CTT GTA CAG CTC GTC CAT GC-3’; the underscores represent the EcoRV and NheI sites added to the primers). The EcoRV-NheI fragment (743 bp) was inserted into the corresponding sites of the pMTWG vector.

In the homomeric interaction assay in S2 cells (Fig. 5, D and E), the following ΔC Spaid plasmids were used: pMT-ΔC Spaid-EGFP.3xFLAG; pMT-ΔC Spaid-3xHA; and pMT-ΔC2 Spaid-EGFP.3xFLAG. To construct the ΔC Spaid-EGFP.3xFLAG plasmid, the pMT-ΔC Spaid-EGFP plasmid was digested with NcoI and NheI to swap the EGFP tag with a gBlocks gene fragment encoding the EGFP.3xFLAG tag sequence (IDT). The ΔC Spaid-3xHA plasmid was generated by replacing the EcoRI-SacI fragment (1,473 bp) of the pMT-ΔC Spaid.358A-3xHA plasmid with the corresponding fragment from pMT-ΔC Spaid-EGFP. To construct the ΔC2 Spaid (1,425 bp) plasmid, a PCR fragment containing a 525-bp portion of *ΔC2 spaid* (nucleotide position 901-1,425, containing an EcoRI site) was amplified from the pMT-ΔC Spaid-EGFP plasmid (forward primer 5’-CTG CAA GTG CGC AAC AAT ATC-3’, reverse primer 5’-CAC <u>GAG CTC</u> ACC ACT TTG TAC AAG AAA GCT GGG TCA TTG ATC TCG TTG ATC ATG CG-3’; the underscore represents the SacI site added to the primer). The EcoRI-SacI fragment (531 bp) was inserted into the pMT-ΔC Spaid-EGFP.3xFLAG plasmid digested with corresponding restriction enzymes. To make a control GFP plasmid (pMT-EGFP.3xFLAG), the pMT-ΔC Spaid-EGFP.3xFLAG plasmid was digested with EcoRV and SacI. After blunting the SacI site by the Quick Blunting Kit (NEB), the digested plasmid was self-ligated.

PrimeSTAR Max DNA Polymerase (Takara Bio) and Mighty Mix DNA ligation kit (Takara Bio) were used for PCR and DNA ligation reactions, respectively.

### Staining and imaging

Salivary glands were dissected out from wandering third instar larvae and fixed in PBS with 4% paraformaldehyde (EM Grade; Electron Microscopy Sciences, 15710) at room temperature for 20 min with gentle rocking. After washing with PBT (PBS containing 0.1% Triton X-100), tissues were treated with a blocking buffer [PBT containing 1% bovine serum albumin (BSA, heat shock fraction; Sigma-Aldrich, A7906)] for 30 min, and incubated with primary antibodies at 4 °C overnight, washed three times in PBT, and incubated with secondary antibodies at room temperature for 90 min. Antibodies were diluted in blocking buffer. For GFP, raw fluorescent signals without antibody staining were detected.

The following primary antibodies were used: rabbit anti-acetyl-Histone H4 (Lys16) (H4K16ac, 1:2,000; Upstate, 07-329); mouse anti-γ-H2Av UNC93-5.2.1 (pH2Av, 1:500; Developmental Studies Hybridoma Bank). The following secondary antibodies were used at a 1:2,000 dilution: donkey anti-mouse IgG Alexa Fluor Plus 555 conjugate (Thermo Fisher Scientific, A32773), donkey anti-rabbit IgG Alexa Fluor Plus 647 conjugate (Thermo Fisher Scientific, A32795). DNA staining was carried out with DAPI (0.5 µg/ml, Nacalai tesque, 19178-91) together with secondary antibody staining.

Stained salivary glands were washed three times in PBT, mounted in ProLong Glass Antifade Mountant (Thermo Fisher Scientific, P36980) and observed under a FV3000 confocal laser scanning microscope (Evident). Images were acquired using a 40x oil immersion objective (UPLXAPO40XO, NA 1.4) with 2x zoom scan (1,024 × 1,024 frame size; about 20-30 sections in 0.42 µm optimal intervals). Transfected S2 cells cultured in plastic multi-well plates were directly observed under the FV3000 confocal microscope with a 10x objective (UPLXAPO10X, NA 0.4). Images were acquired at six areas per well (800 × 800 frame size; 11 sections in 6.0 µm intervals) for quantitative analysis.

### Image analysis and processing

Two confocal z-sections of salivary gland cells were selected and max-projected in Fig. 3 and S3. All confocal z-sections of S2 cells were max-projected in Fig. 4A, 5A, and S5A. The brightness and contrast of the presented images were adjusted by Fiji software (Fiji Is Just ImageJ) (*69*). The adjustment was performed uniformly on the entire images.

For the relative quantification of GFP signals of transfected S2 cells (Fig. 4B, 5B, and S5B), all z-sections were max-projected by a custom macro with Fiji software. Image analysis was performed by custom R scripts with the EBImage package (*70*). The maximum projection images of GFP and mCherry were smoothed by a Gaussian filter and binarized by the moving average method. GFP and mCherry signals were measured by image integration, and the value of GFP was divided by that of mCherry to calculate the relative fluorescence of GFP.

### Biochemical assays

For the ubiquitination assay, transfected S2 cells were pelleted at 1,000 g and rinsed twice with ice-cold PBS. The cells were lysed in 200 µl RIPA buffer (50 mM Tris-HCl pH 7.6, 150 mM NaCl, 1% NP-40 substitute, 0.5% sodium deoxycholate, 0.1% SDS; Nacalai tesque, 16488-34) containing 1 mM EDTA, 5 mM N-ethylmaleimide (NEM, Nacalai tesque, 15512-24), 40 µM PR-619 (LifeSensors, SI9619), 1x cOmplete ULTRA (Roche, 5892791001), and 1 mM Pefabloc SC (Roche, 11429868001). After a 15 min incubation on ice with periodic mixing, the samples were pulse sonicated, then incubated on ice for another 15 min. Cell lysates were cleared by centrifugation at 13,000 g at 4 °C for 10 min and the supernatant was collected and stored at -80 °C until use. Protein concentration was quantified using the BCA protein assay kit (Takara Bio, T9300A). Cell lysates (about 200 µl) were diluted 2-fold with dilution buffer (50 mM Tris-HCl pH 7.5, 150 mM NaCl, 0.5 mM EDTA) supplemented with 5 mM NEM, 40 µM PR-619, 1x cOmplete ULTRA, and 1 mM Pefabloc SC. For immunoprecipitation, 6 µl GFP-Trap magnetic agarose (ChromoTek, gtmak-20) pre-equilibrated with dilution buffer was added to the diluted lysates, then gently rotated at 4 °C for 2 h. Magnetically separated beads were washed once with dilution buffer, three times with stringent washing buffer (8M Urea, 1% SDS in PBS, prepared just before use), and once with 1% SDS in PBS (*44, 45*). Bound proteins were eluted with 2x sample buffer (Bio-Rad, 1610747) containing 0.1M TCEP-HCl (Nacalai tesque, 07277-61), incubated at 70 °C for 30 min, then resolved by SDS-PAGE on 4-15% precast gradient gels (Mini-PROTEAN TGX Gels, Bio-Rad, 4561084) in Tris/Glycine/SDS buffer (Bio-Rad, 1610732) for 30 min with 200V constant voltage. Proteins were transferred to 0.2 µm PVDF membranes (Trans-Blot Turbo Transfer Pack, Bio-Rad, 1704156) using the “High MW” setting of the Trans-Blot Turbo Transfer System (Bio-Rad). Blotted membranes were rinsed briefly with TBS-T [TBS (20 mM Tris-HCl pH 7.6, 150 mM NaCl) containing 0.1% Tween 20], treated with a blocking buffer (TBS-T containing 1% BSA) at room temperature for 30 min, or Bullet Blocking One (Nacalai tesque, 13779-14) at room temperature for 5 min, and incubated with primary antibodies at 4 °C overnight; the membranes were then washed three times in TBS-T and incubated with peroxidase-conjugated secondary antibodies at room temperature for 1 h. Primary and secondary antibodies were diluted in the blocking buffer. After washing three times with TBS-T, the membranes were incubated with ECL Prime Western Blotting Detection Reagent (Amersham, RPN2232) at room temperature for five min, and chemiluminescent signals were detected with a ChemiDoc Imaging System (Bio-Rad).

For the co-immunoprecipitation assay, cell lysates were prepared as indicated above except that cells were lysed with RIPA buffer without SDS (Nacalai tesque, 08714-04) and lysates were passed through a 27G needle 10 times; the volume was then adjusted to 400 µl with dilution buffer prior to immunoprecipitation using GFP-Trap magnetic agarose beads. After protein binding, the beads were washed three times with lysis buffer (50 mM Tris-HCl pH 7.5, 150 mM NaCl, 0.5% Triton X-100, 0.5 mM EDTA) containing 1x cOmplete ULTRA and 1 mM Pefabloc SC, and bound proteins were analyzed by western blotting.

The following antibodies were used: mouse anti-Ubiquitin P4D1 (1/1,000; Santa Cruz, sc-8017); rabbit anti-GFP (1/10,000; Invitrogen, A-11122); mouse anti-FLAG M2 (1/10,000; Sigma-Aldrich, F1804); rat anti-HA 3F10 (1/5,000; Roche, 11867423001); mouse anti-β-Actin C4 (1/5,000; Santa Cruz, sc-47778). The following peroxidase-conjugated secondary antibodies were used (1/10,000; purchased from Jackson ImmunoResearch): donkey anti-mouse IgG (715-035-150); donkey anti-rabbit IgG (711-035-152); donkey anti-rat IgG (712-035-153).

### Protein domain search, multiple alignment, and 3D modeling

The amino acid sequence of FL Spaid was analyzed by the InterPro website (*39*) with default parameters. The following three predictions associated with OTU/cysteine protease were obtained: SSF54001 (“Cysteine proteinases”, amino acids 264-461, e-value: 1.18E-15), PF02338 (“OTU-like cysteine protease”, amino acids 344-438, e-value: 6.50E-06), and PS50802 (“OTU domain profile”, amino acids 279-465, score: 13.786528). In this paper, the last prediction was adopted according to the results of the OTU domain alignment shown in Fig. S1A.

For the multiple alignment of the OTU domain sequence, Jalview v2.11.2.4 was used. I also referred to the alignment presented in ref. (*27*). Protein sequences of OTU family proteins from *Homo sapiens* (Hsap), *Drosophila melanogaster* (Dmel), *Chlamydia pneumoniae* (Cpne), *Legionella pneumophila* (Lpne; only the N-terminal OTU domain sequence was used) were obtained from UniProtKB (accession numbers in parentheses): OTU1_Hsap (Q5VVQ6); OTUB1_Hsap (Q96FW1); OTUB2_Hsap (Q96DC9); OTUD1_Hsap (Q5VV17); OTUD3_Hsap (Q5T2D3); OTUD4_Hsap (Q01804); OTUD5_Hsap (Q96G74); OTU6A_Hsap (Q7L8S5); OTU6B_Hsap (Q8N6M0); OTU7A_Hsap (Q8TE49); OTU7B_Hsap (Q6GQQ9); TNAP3_Hsap (P21580); VCIP1_Hsap (Q96JH7); ZRAN1_Hsap (Q9UGI0; a.k.a. TRABID); ALG13_Hsap (Q9NP73); OTUL_Hsap (Q96BN8); OTU_Dmel (P10383); OTU1_Dmel (Q9VRJ9); TRBID_Dmel (Q9VH90); ChlaOTU_Cpne (Q9Z868); LotA_Lpne (Q5ZTB4).

3D models of the OTU domain of Spaid (279-465 aa) were generated by the Phyre2 server using the intensive mode (*40*) (Fig. S1B and Table S1, sheet1) and ColabFold software (*41*) powered by the AlphaFold2 program (*42*) (Fig. S1C), respectively. Structural comparison of the ColabFold model was performed by the DALI server (*43*) (Table S1, sheet2). Default parameters were utilized in all programs. Obtained 3D models were visualized by PyMOL v2.5.0 (Schrödinger).

### Statistical analysis

R software v4.0.3 (the R Foundation) was used for all statistical analyses. Multiple comparisons in Fig 2A, 4B, 5B, S2A, and S5B were performed using the Steel-Dwass test by the pSDCFlig function in the NSM3 R package (*71*). The Mann-Whitney U test (two-tailed) was used in Fig 2B and S2B. *P*-values less than 0.05 were considered as significant. Exact *P*-values are listed in Table S2.

## ACKNOWLEDGMENTS

I thank the Bloomington *Drosophila* Stock Center in the USA and the Department of *Drosophila* Genomics and Genetic Resources at Kyoto Institute of Technology in Japan for fly stocks, and the Developmental Studies Hybridoma Bank at the University of Iowa in the USA for monoclonal antibodies. I also thank Catherine Regnard and Peter Becker for providing plasmid vectors, Masayuki Miura for the cell stock, Makoto Hayashi for comments and suggestions on the work, James Alan Hejna for comments and English editing, Natsuya Oura and the members of Tadashi Uemura’s laboratory for discussion and support.

## Author contributions

Conceptualization: TH

Methodology: TH

Investigation: TH

Visualization: TH

Supervision: TH

Writing—original draft: TH

Writing—review & editing: TH

## Data and materials availability

Correspondence and requests for materials should be addressed to Toshiyuki Harumoto. All data are available in the main text or the supplementary materials.

## SUPPLEMENTARY MATERIALS

**Fig. S1.**
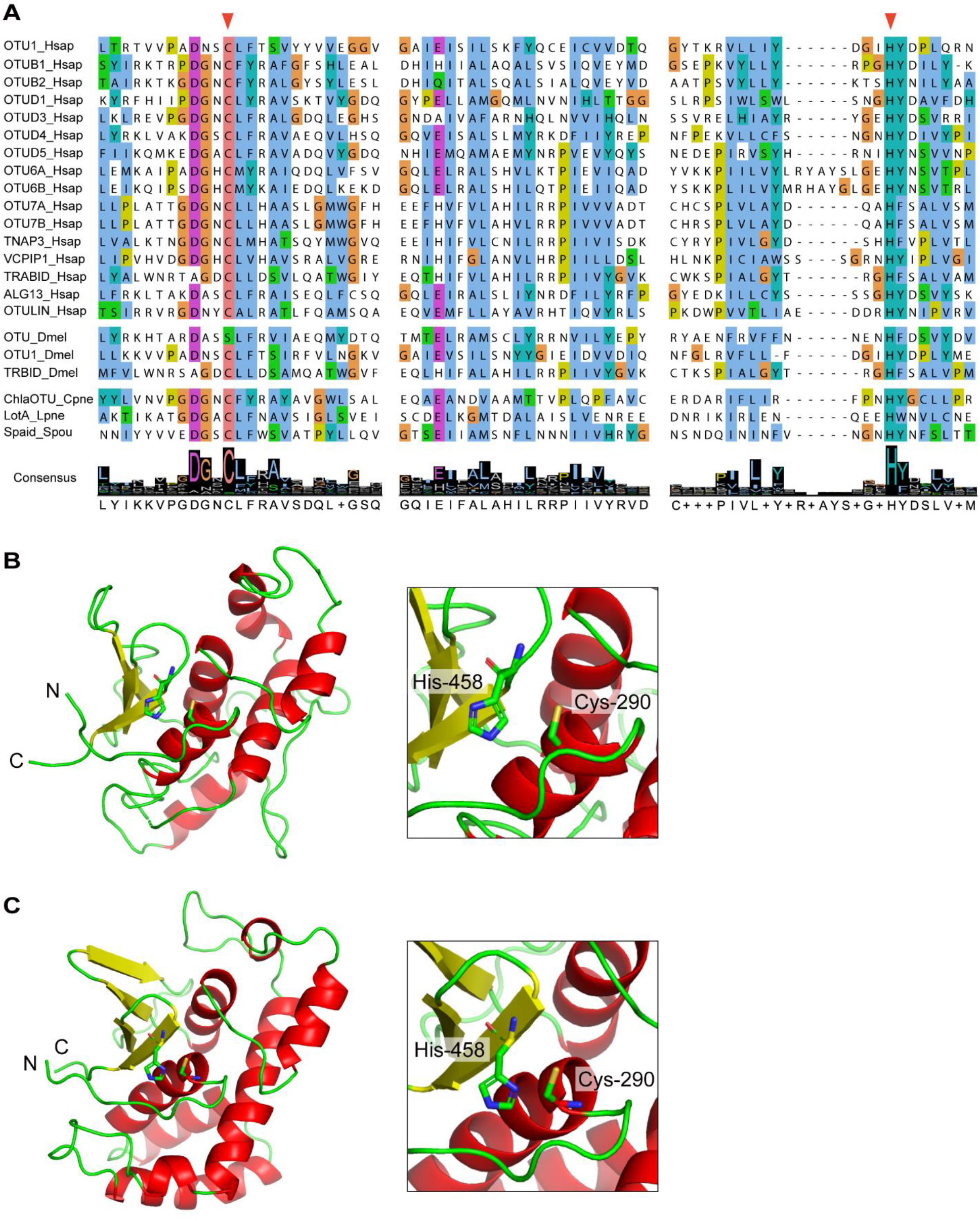
Multiple alignment and 3D structure of the OTU deubiquitinase domain of Spaid. (**A**) The predicted OTU domain of Spaid (amino acid positions 279-465, bottom row) partially aligned with representative OTU family proteins conserved in eukaryotes (*Homo sapiens*, Hsap; *D. melanogaster*, Dmel) and bacteria (*Chlamydia pneumoniae*, Cpne; *Legionella pneumophila*, Lpne; *S. poulsonii*, Spou) (see Materials and Methods for details). Red triangles at the top represent a catalytic dyad composed of cysteine (C) and histidine (H), which corresponds to Cys-290/His-458 in the predicted OTU sequence of Spaid. The consensus sequence logo is indicated at the bottom. (**B, C**) Predicted 3D models of the OTU domain of Spaid, generated by Phyre2 (B) and ColabFold (C), respectively. Secondary structural motifs (helix, sheet, loop) are colored differently. Side chains of Cys-290 and His-458 are shown. Close-up images of the active site region comprising the Cys-290/His-458 dyad are indicated in the right boxes. N, N-terminal; C, C-terminal.

**Fig. S2.**
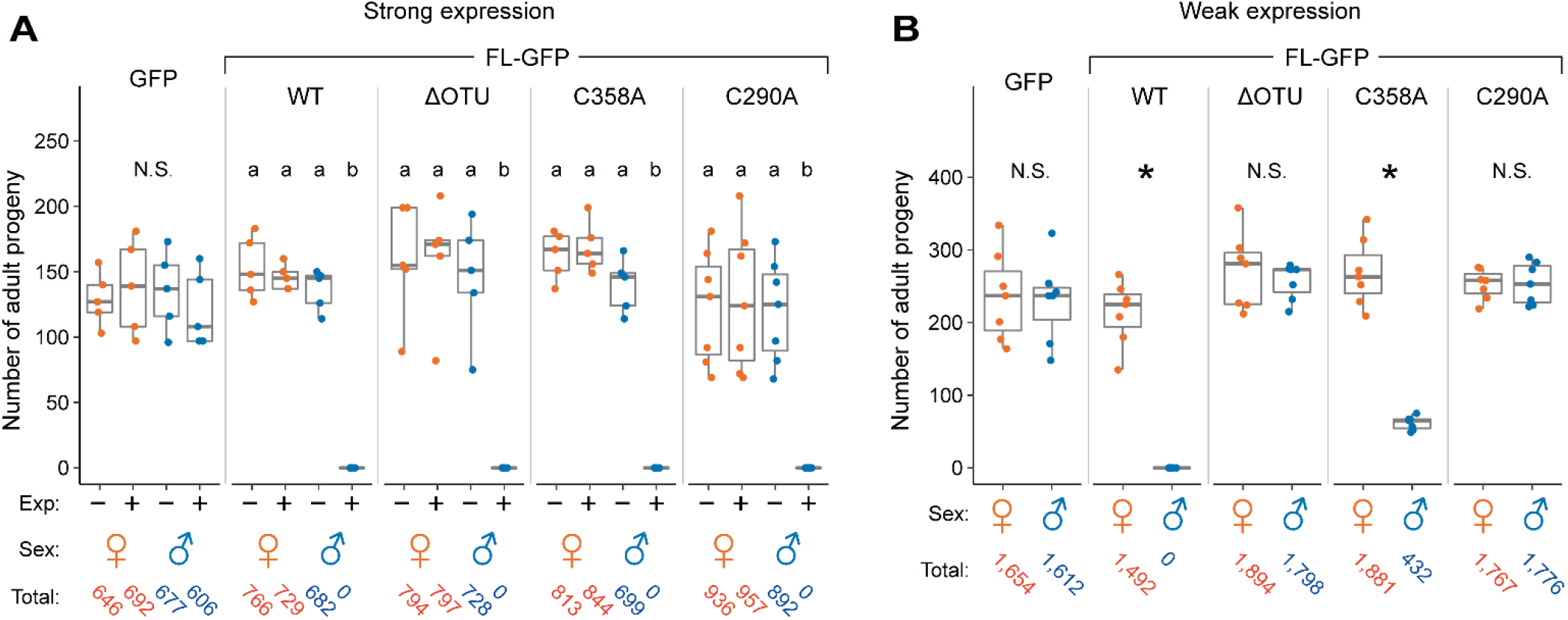
OTU-deficiency attenuates the male-killing activity of FL Spaid. *UAS-GFP* (negative control), *UAS-FL Spaid* (WT) and its derivative lines (ΔOTU, C358A, and C290A) were crossed with the *actin-GAL4* (**A**, strong expression; n = 5 independent crosses, n = 7 for C290A) and *armadillo-GAL4* (**B**, weak expression; n = 7 independent crosses) driver lines, respectively. The numbers of female (red) and male (blue) progeny obtained from the crosses are shown. In **A**, offspring with both *GAL4* and *UAS* (+) or with only *UAS* (–) were counted separately. Different letters (Steel–Dwass test, **A**) and asterisks (two-tailed Mann–Whitney U test, **B**) represent statistically significant differences (P < 0.05). NS, not significant (P > 0.05) (See also Table S2). Box plots indicate the median (bold line), upper and lower quartiles (box edges), and the maximum/minimum values (whiskers). Dot plots show individual data points. The total counts for each genotype and sex are shown at the bottom.

**Fig. S3.**
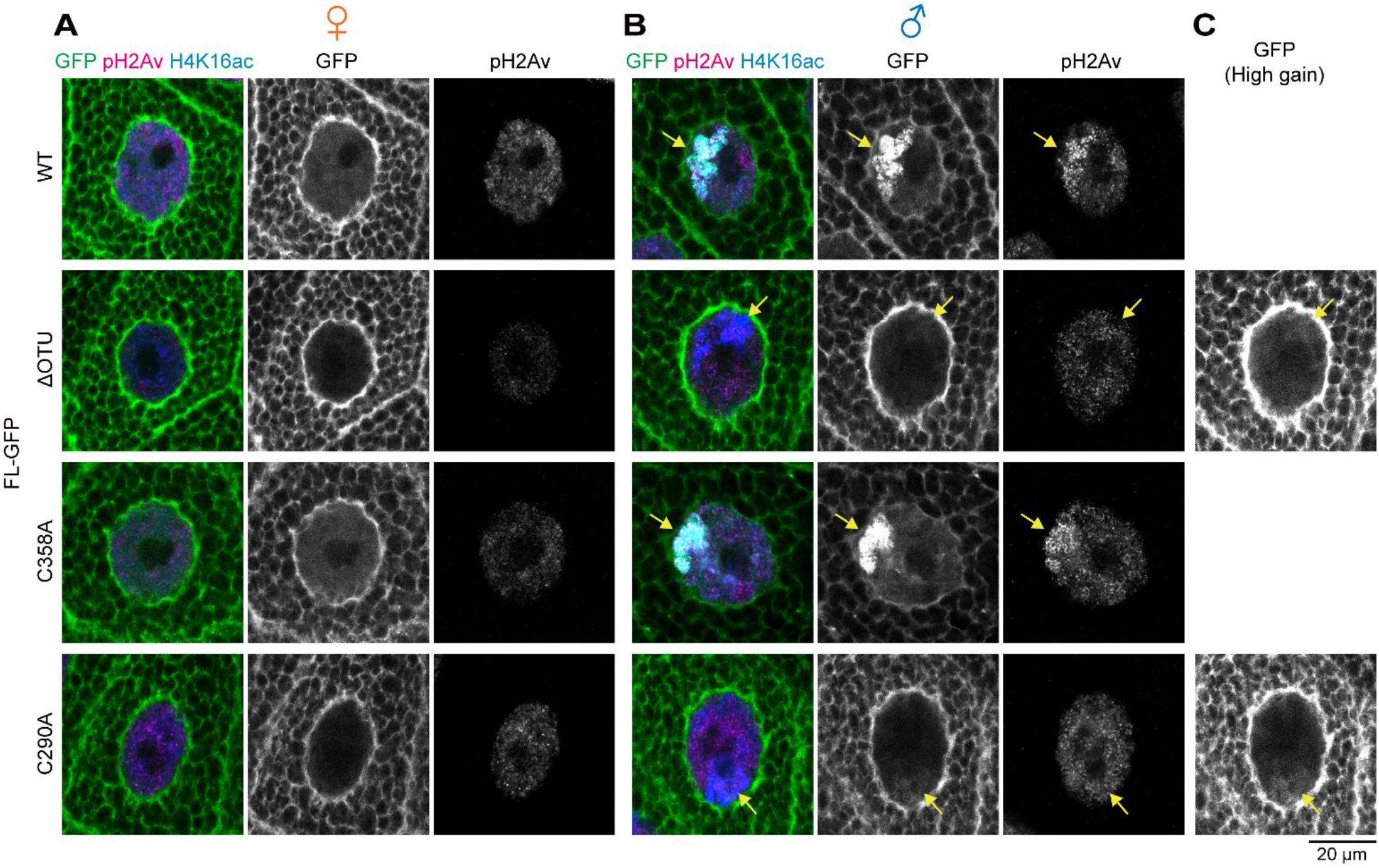
Subcellular localization of FL Spaid. Female (**A**) and male (**B**) larval salivary gland cells expressing FL Spaid-GFP (green). WT (female, n = 8; male, n = 8), ΔOTU (female, n = 12; male, n = 11), C358A (female, n = 11; male, n = 10), and C290A (female, n = 7; male, n = 12) derivatives were immunostained for DNA damage (pH2Av, magenta) and histone H4 lysine 16 acetylation (H4K16ac, blue). Yellow arrows represent the accumulation of FL Spaid-GFP and pH2Av on the male X chromosome labeled with the dosage compensation machinery (marked with strong H4K16ac signals). (**C**) High-gain images of ΔOTU and C290A reveal faint GFP signals on the male X chromosome.

**Fig. S4.**
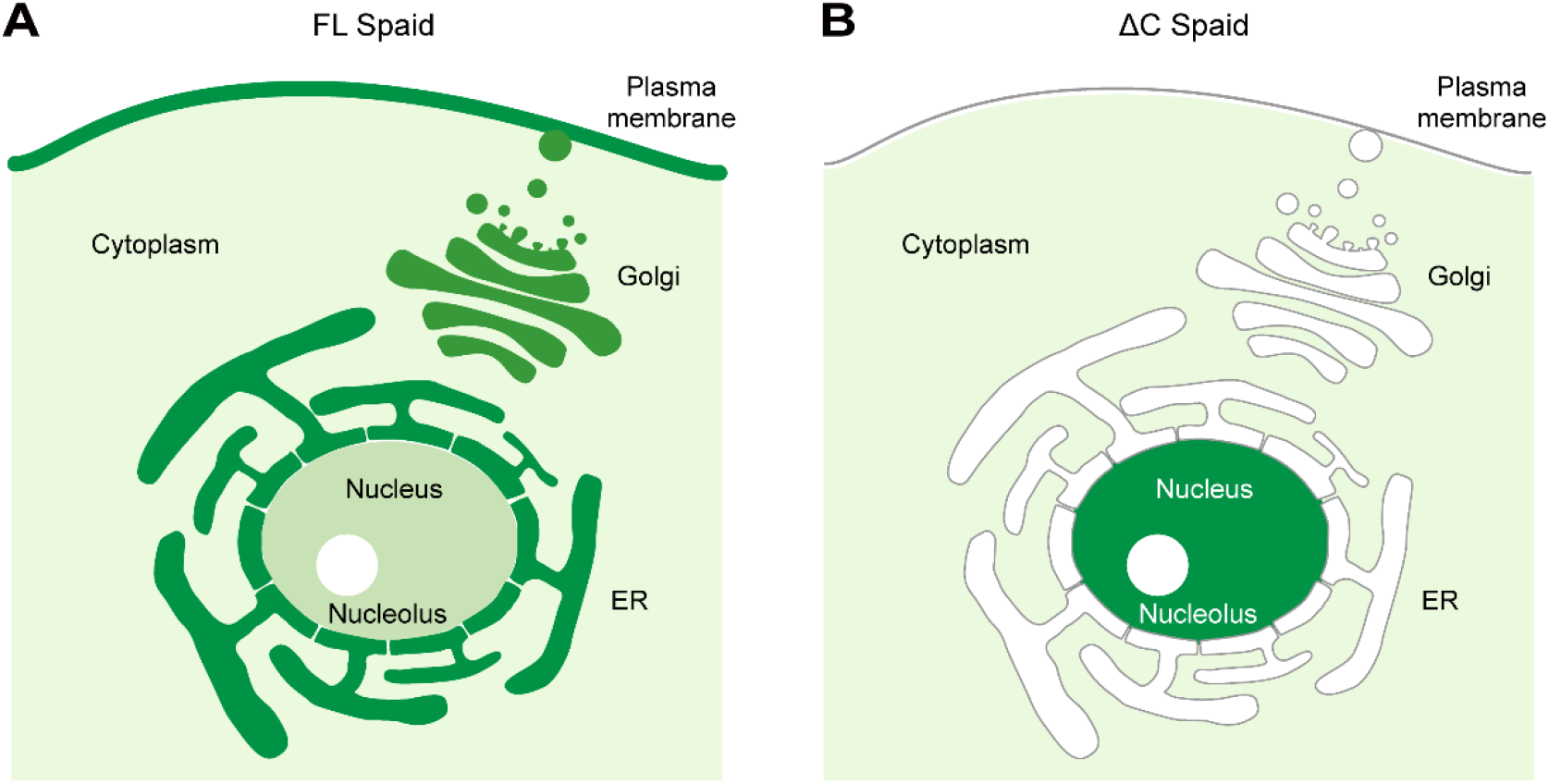
Schematics of intracellular localization of FL and ΔC Spaid. When artificially expressed in the host cells, (**A**) a substantial amount of FL Spaid is delivered to the intracellular membrane system including the endoplasmic reticulum (ER), Golgi apparatus, and plasma membrane (dark green in **A**) with the help of its hydrophobic region (HR) (corresponding to the images shown in Fig. S3). Meanwhile, (**B**) ΔC Spaid, which lacks the C-terminal region containing HR, is predominantly accumulated within the nucleus (dark green in **B**) (corresponds to the images shown in Fig. 3).

**Fig. S5.**
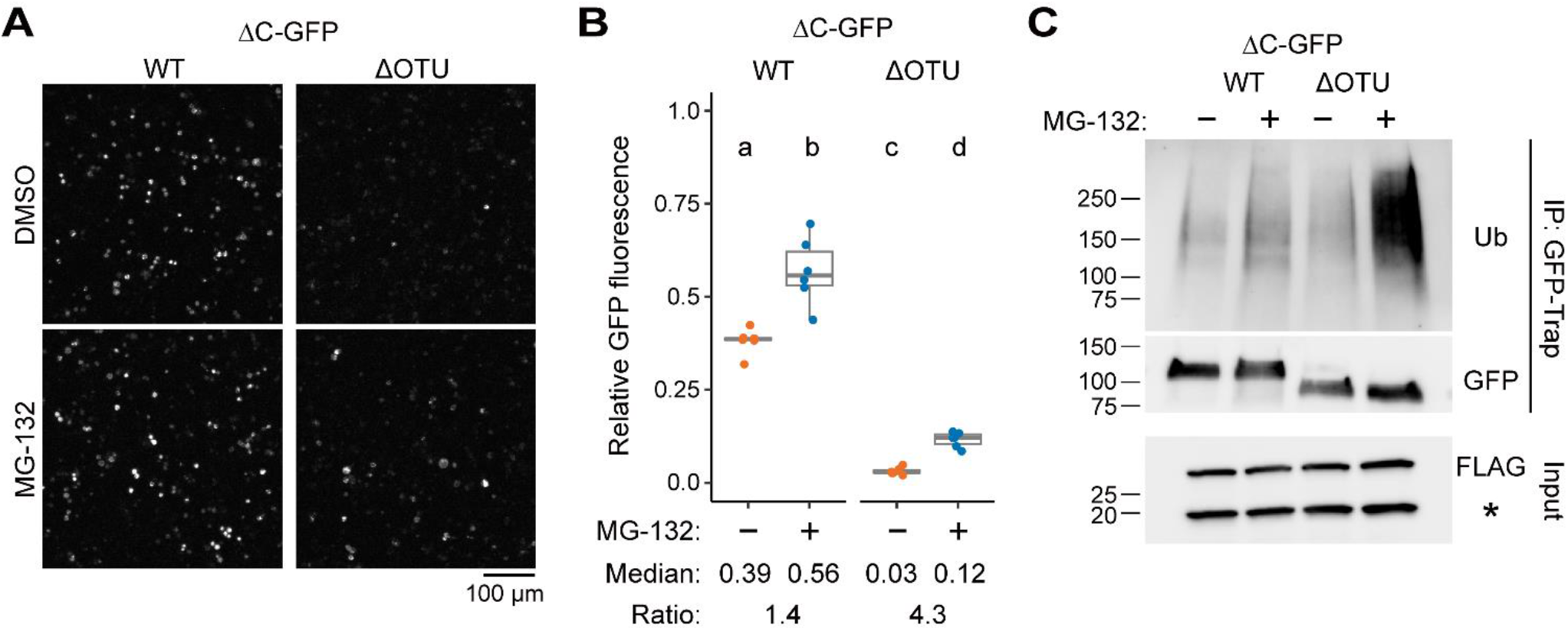
The OTU domain of Spaid stabilizes itself by counteracting the host ubiquitin-proteasome pathway. (**A**) S2 cells expressing ΔC Spaid-GFP. The raw GFP signals of WT and ΔOTU derivatives are indicated. Cells were treated with DMSO (top) or proteasome inhibitor MG-132 (bottom) for 9 h. (**B**) The relative GFP fluorescence of WT and ΔOTU derivatives of ΔC Spaid-GFP expressed in S2 cells cultured without or with MG-132. The same data sets used in (**A**) were analyzed (n = 6 areas per well in each condition). Other explanations are the same as in Fig. 4B. Different letters (Steel–Dwass test) represent statistically significant differences (P < 0.05) (See also Table S2). (**C**) Ubiquitination assay using GFP-Trap. WT and ΔOTU derivatives of ΔC Spaid-GFP were immunoprecipitated (IP) from S2 cells cultured without or with MG-132. Other explanations are the same as in Fig. 4C.

**Table S1. The results of the 3D structural modeling of Spaid OTU.**(**Sheet1**) The output of the Phyre2 server search. The top 40 templates used for the 3D model prediction of the Spaid OTU sequence (279-465 aa) are listed. Confidence of the modeling (100% maximum, shown in a red-white scale), sequence identity (% i.d.) between Spaid OTU, Protein Data Bank (PDB) IDs and associated information, names of molecules and source organisms are indicated. vOTU, viral OTU. The Phyre2 server selected six templates (colored in yellow) to generate the final 3D model indicated in Fig. S1B. (**Sheet2**) The output of the DALI server search. The top 20 proteins with high structural similarities between the ColabFold model of Spaid OTU are listed. Z scores (optimized similarity score, shown in the red-white scale), RMSD (root-mean-square deviation), sequence identity (% i.d.) between Spaid OTU, PDB IDs and associated titles, names of molecules, and source organisms are indicated.

**Table S2.The exact *P*-values of statistical analyses.***P*-values less than 0.05 (colored in red) were considered significant.

